# Interaction between DNA damage response, translation and apoptosome determines cancer susceptibility to TOP2 poisons

**DOI:** 10.1101/614024

**Authors:** Chidiebere U Awah, Li Chen, Mukesh Bansal, Aayushi Mahajan, Jan Winter, Meeki Lad, Louisa Warnke, Edgar Gonzalez-Buendia, Cheol Park, Zhang Daniel, Eric Feldstein, Dou Yu, Markella Zannikou, Irina V. Balyasnikova, Regina Martuscello, Silvana Konerman, Balázs Győrffy, Kirsten B Burdett, Denise M Scholtens, Roger Stupp, Atique Ahmed, Patrick Hsu, Adam Sonabend

## Abstract

Topoisomerase II poisons are one of the most common class of chemotherapeutics used in cancer. We show that glioblastoma (GBM), the most malignant of all primary brain tumors in adults is responsive to TOP2 poisons. To identify genes that confer susceptibility to this drug in gliomas, we performed a genome-scale CRISPR knockout screen with etoposide. Genes involved in protein synthesis and DNA damage were implicated in etoposide susceptibility. To define potential biomarkers for TOP2 poisons, CRISPR hits were overlapped with genes whose expression correlates with susceptibility to this drug across glioma cell lines, revealing ribosomal protein subunit RPS11, 16, 18 as putative biomarkers for response to TOP2 poisons. Loss of RPS11 impaired the induction of pro-apoptotic gene APAF1 following etoposide treatment, and led to resistance to this drug and doxorubicin. The expression of these ribosomal subunits was also associated with susceptibility to TOP2 poisons across cell lines from multiple cancers.

**Graphical Abstract:** 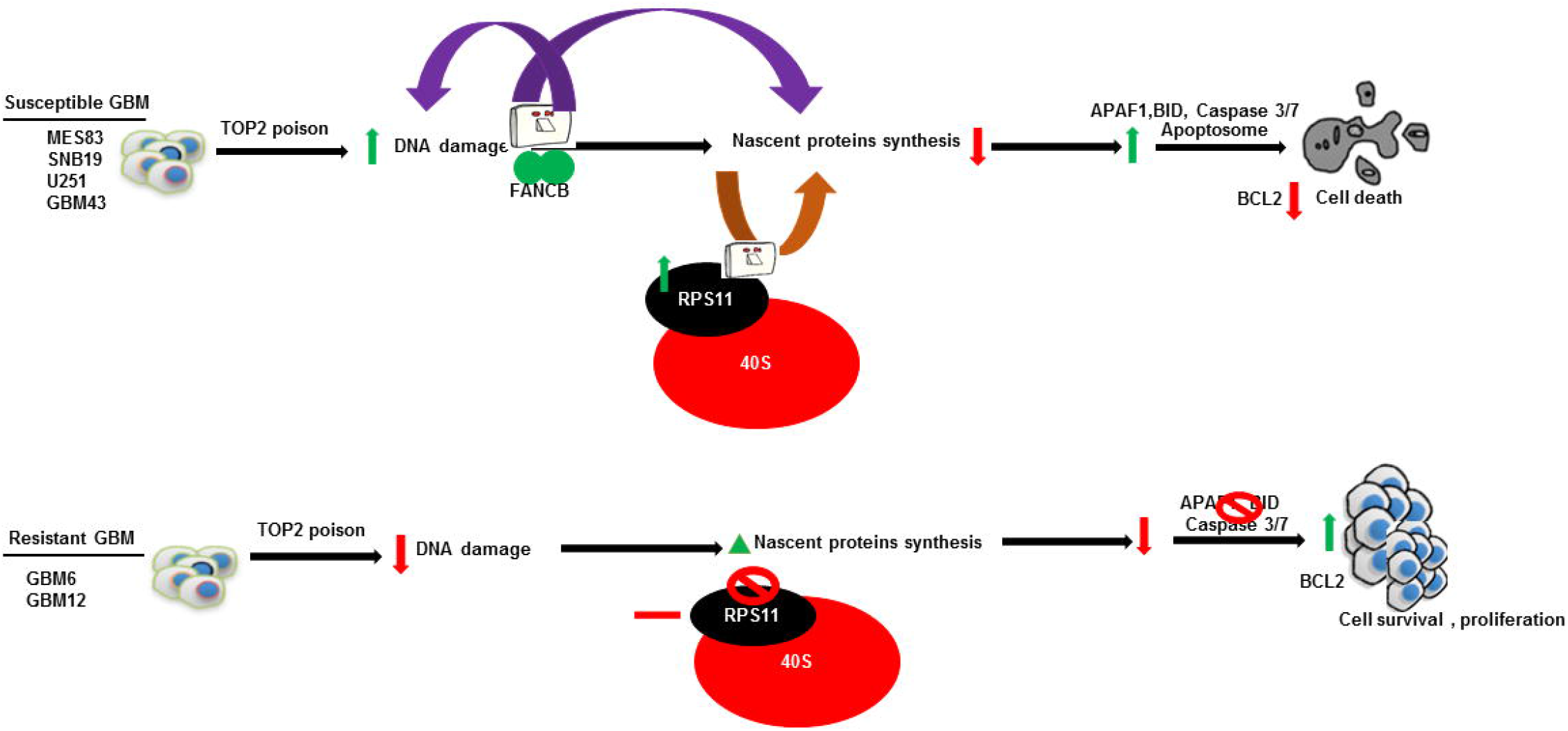

## Introduction

Glioblastoma (GBM) remains the most lethal of all primary brain tumors in adults. The standard therapy for this disease include maximal surgical resection, radiation therapy, chemotherapy with the alkylating agent temozolomide, and more recently, the use of tumor-treating electrical field therapy. Despite this multi-modal therapy, the median survival is approximately 2 years^1^. Such a uniform therapeutic approach contrasts with the molecular diversity of this disease. GBM are notorious for their unpredictable response to therapies, which ultimately contributes to the poor prognosis. To characterize this complexity, several iterations of molecular classifications have been performed based on gene expression patterns, genetic alterations, and DNA methylation^2, 3, 4^ In this context, major efforts are focused on utilizing gene expression patterns to predict unique tumoral vulnerability and inform the choice of specific drugs for individual patients.

Topoisomerase II (TOP2) are enzymes are molecular machines that unwinds DNA during replication and transcription to relax the torsional stress of DNA folding. TOP2 poisons etoposide and doxorubicin, which induce double-strand DNA breaks, are widely used for different cancers^5, 6^ Etoposide is typically used for testicular cancer and small cell lung cancer as these tumors are considered susceptible to this drug. Whereas etoposide and doxorubicin are not commonly used for gliomas, these drugs are also effective in an elusive subset of these tumors^7, 8^ Clinical trials in recurrent gliomas show that some patients responded to etoposide-containing regimens, and this response also led to a survival benefit ^9, 10, 11^. We previously showed that some human glioma cell lines are as susceptible to etoposide as testicular cancer cell lines (the most susceptible cancer to this drug), suggesting that the histological diagnosis might be less important than individual tumor biology predicting response to this drug^7^. In this context, the criteria and molecular signatures for patient selection remains a major challenge for effective therapy using TOP2 poisons for cancer and gliomas in particular, as there are no reliable biomarkers for these drugs.

Rapid advances of the CRISPR-Cas9 genome editing technology have allowed unbiased interrogation of the mammalian genome and efficient linking of genotype and function^12, 13^. CRISPR-based knock-out (KO) screening libraries have been optimized to maximize on-target gene editing. Through the introduction of 3-4 independent sgRNAs per gene, functional consequences resulting from gene inactivation can be assessed, minimizing false-positive results from off-target KO^12, 13, 14, 15^. Taking advantage of this technology to investigate the molecular mechanisms involved in glioma susceptibility to etoposide, we performed a genome scale CRISPR KO screen in cells undergoing treatment with this drug. In order to discover a biomarker for personalizing this therapy for GBM, we have overlapped the genes that conferred etoposide susceptibility in our CRISPR screen, with genes whose expression is associated with susceptibility to this drug across glioma cell lines. This approach led to a short list of biomarker candidates that are experimentally implicated and correlatively associated with susceptibility to this drug. Our results show that ribosomal subunit proteins (RPS11, RPS16 and RPS18) influence glioma susceptibility to TOP2 poisons, and that the expression of these genes is associated with response to TOP2 poisons across cell lines from multiple cancers. We found that RPS11 modulates the expression of pro-apoptotic protein APAF1, which is upregulated following etoposide treatment, and is required for cell death from this therapy. In brief, our results suggest that protein synthesis, DNA damage and apoptosis influence susceptibility to etoposide across GBMs and introduce RPS11 as a promising biomarker for response to TOP2 poisons.

## Results

### Translation-related genes and DNA damage repair pathways confer glioma susceptibility to etoposide

We performed a genome-wide scale CRISPR KO screen using a clinically relevant dose of etoposide (5 μM) to identify genes that influence human glioma susceptibility to etoposide. Several clinical studies quantified intratumoral concentrations of etoposide in gliomas and brain metastases following systemic administration of this drug and found an intratumoral concentration range between 2-6 μM in the tumor tissue [16-18]. Moreover, the treatment with etoposide at 5 μM for 72 hrs led to 80% cells death in susceptible glioma cell line SNB19, whereas resistant cell lines showed minimal cell death (Fig **1a**). Thus, our CRISPR KO screen experiment was performed selecting with etoposide 5 μM or DMSO, etoposide solvent, for 14 days (Fig **1b**). This treatment led to strong selection with less than 1% of cells surviving treatment (Supplementary Fig **1A**). The cumulative frequency of sgRNA sequencing reads showed that the guides from etoposide were distinct from the counts obtained by sequencing the library plasmid (library) and those from Day 0 (post-puromycin selected cells) (Supplementary Fig **1B**). We repeated the etoposide arm of the CRISPR experiment and validated the reproducibility of our CRISPR screen for most targets with both screens showing Spearman’s r^2^=0.68 (p<0.0001) both on the gene and sgRNA level (Supplementary List **1A**). This screen showed that sgRNAs for the genes involved in ribosome and protein synthesis were over-represented in the etoposide as opposed to DMSO-treated cells (p<1.0E^-6^, Fisher’s exact test with Benjamini multiple hypothesis correction) (Supplementary Fig **1C**).

**Fig 1:**
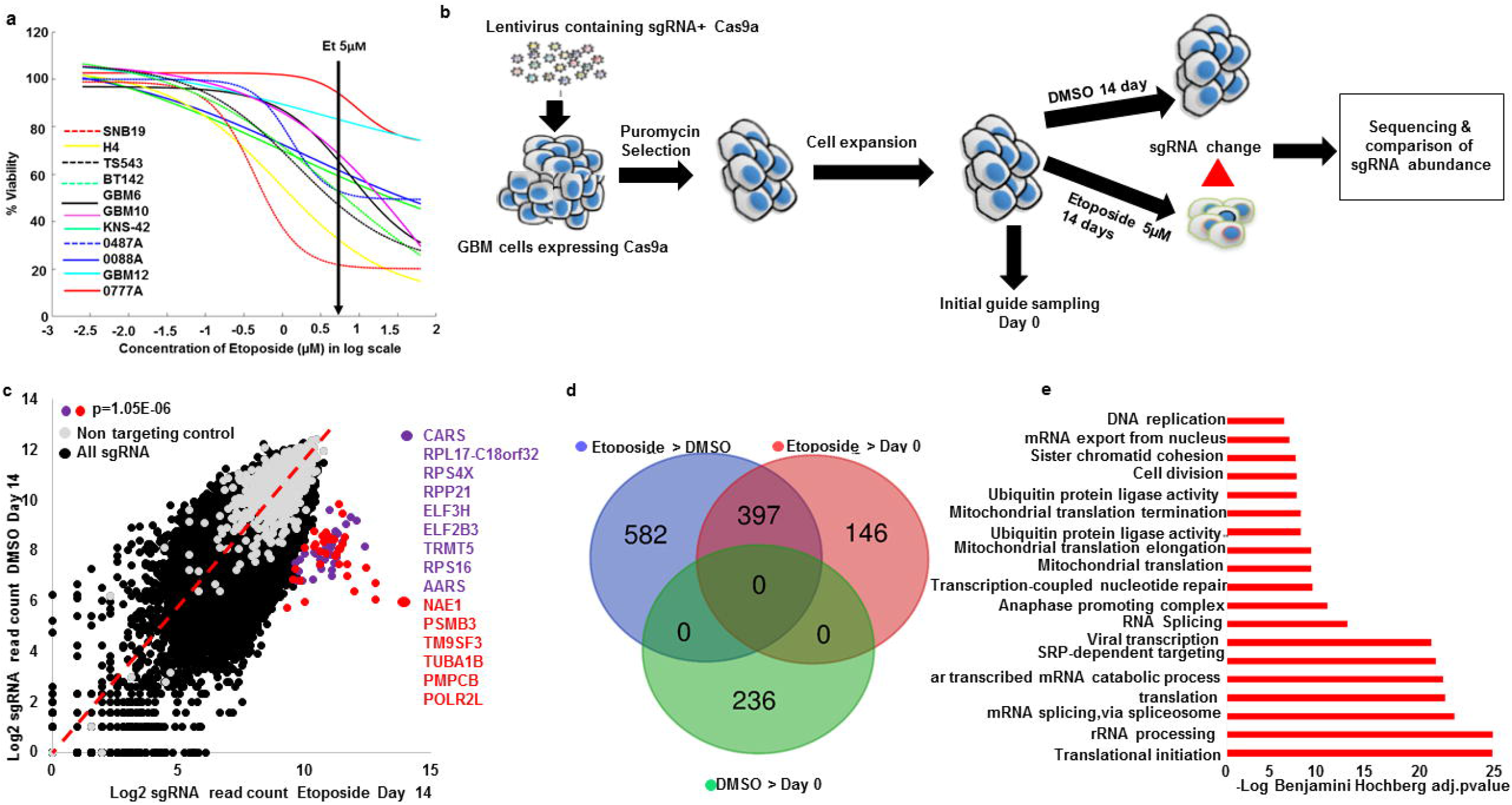
CRISPR screen in glioma cells reveals genes that confer susceptibility to etoposide. **(a)** Etoposide dose response curves of 11 human glioma cell lines treated with etoposide (2-40μM) and DMSO for 72hrs. **(b)** Schematic depiction of the CRISPR screen experiment performed in this study. Vector containing 76,441 sgRNA library expressing Cas9a were packaged into lentivirus, which was spinfected into SNB19 cells. The transfected cells were selected with puromycin for 96hrs. Cells were expanded and split into etoposide treatment and DMSO for 14 days with 1×10^8^ cells per condition (1298X coverage). Unique barcoded primers were used to amplify the library, puromycin/Day 0, DMSO and etoposide selected guides. These samples were pooled and sequenced. sgRNA enrichment was analyzed using CRISPRAnalyzer ^45^. All experiment was done in two independent replicates except for DMSO that was performed once. **(c)** Scatter plot depicts the genes with highest sgRNA enrichment by etoposide (p<0.000001). Ribosomal and tRNA synthetase genes marked in purple. Red genes represent those related to ubiquitin, proteasome, tubulin and RNA Polymerase II subunits. The rest of sgRNA are represented in black and the non-targeting controls in gray. **(d)** Venn diagram show number of genes that were enriched in comparisons between different experimental conditions. For this p<0.01 was used as a cutoff for hit calling. **(e)** Bar chart shows the enriched gene ontology themes from the 397 genes whose KO enrichment overlapped between etoposide>DMSO and etoposide >puromycin. Benjamini Hochberg adjusted pvalue for gene ontology enrichment cutoff, p=0.001.

To identify the pathways implicated in glioma susceptibility to etoposide, we first analyzed genes that were enriched by etoposide compared to DMSO and/or Day 0. Using a cutoff for hit calling by sgRSEA enriched of p<0.01 (Wilcoxon), this analysis showed 979 genes whose KO was uniquely enriched by etoposide compared to DMSO (etoposide>DMSO) (Fig **1d**). 543 genes whose KO were enriched in etoposide compared to Day 0 (etoposide>Day 0) (Fig **1d**). 397 genes whose KO was enriched and overlapped between etoposide >DMSO and in etoposide > Day 0 selected cells (Fig **1d**, Supplementary List **1B**). 236 genes whose KO was enriched in DMSO compared to Day 0, and these genes were not found to be enriched in etoposide >DMSO nor in etoposide > Day 0 comparison (Fig **1d**). We used the 397 genes whose KO showed enrichment and overlapped between etoposide > DMSO and etoposide > Day 0 (Fig **1d**) to perform a gene ontology analysis (DAVID). We found ontology themes related to translation as the most over-represented among the group of 397 genes (Fig **1e**, Supplementary List **1B**, and **2A**). Amongst the genes enriched in translational machinery, the most overrepresented were ribosomal subunit proteins, followed by mitochondrial ribosomal proteins and tRNA synthetase (Supplementary Fig **1C**). Etoposide induces doublestrand DNA breaks^5, 6^ and on the other hand, gliomas are known to exhibit relatively high genome instability^19^. Thus, we sought to investigate whether a component of the DNA damage and repair pathways is required for susceptibility of TOP2 poisons in gliomas. We found that 57 out of 348 genes previously implicated in DNA damage and repair^20^ had their KO clone enriched in Etoposide>DMSO or in Etoposide>Day 0 (Fischer exact test for enrichment p=2.3E^-12^), and 24 genes whose KO was enriched by etoposide compared to DMSO and Day 0 (Fisher exact test for enrichment p=1.9E^-7^) (Supplementary List **2B**). Using the 57 DNA damage and repair genes found to be enriched in etoposide > DMSO or in etoposide > Day 0 sample, we performed a gene ontology analysis, and found the theme of double strand DNA repair via homologous recombination to be enriched (p=2.7E^-6^ Fisher’s exact test with Benjamini multiple hypothesis corrections, Supplementary Fig **2A**), implicating these genes in the known mechanism of TOP2 poisons of inducing double-strand DNA breaks.

Other groups have performed genome-wide KO screens using etoposide in leukemia using CRISPR technology ^20, 21^ and also with other strategies for genome-wide screens to elucidate susceptibility to this drug in cancer^22, 23^. Whereas there are differences in the cancer cell line (e.g., leukemia vs. glioma), drug concentration, exposure period and analysis, our screen validated 20 out of 25 genes hits previously reported by these studies (Fig **1d**, Supplementary List **2B**). In conclusion, the CRISPR screen reveals DNA damage and proteins involved in translation as key regulators of response to TOP2 poison.

### Susceptibility to TOP2 poisons across gliomas is linked to DNA damage

We previously showed that susceptibility to TOP2 poison etoposide varies significantly across human cancer cell lines^7, 8^. To investigate whether individual cancer cell line susceptibility to these drugs relates to the established mechanism of action, we compared the area under the dose response curve (AUC) of individual cell lines for etoposide versus doxorubicin, and versus that of other chemotherapy agents that are not TOP2 poisons (n=665, Cancer Cell Line Encyclopedia)^23,24^. This analysis showed a high correlation of susceptibility to these TOP2 poisons across cancer cell lines, and similar results were observed when the analysis was restricted to gliomas (Fig **2a**). Yet, no correlation was found between either of the TOP2 poisons versus cisplatin or cytarabine, chemotherapeutics with a different mechanism of action. The results of DNA damage response theme represented by our CRISPR hits led us to hypothesize that individual variation in tumor susceptibility to TOP2 poisons relates to the mechanism of action for these drugs. To investigate whether differential etoposide susceptibility relates to DNA damage response, we quantified yH2AX (phospho gamma H2AX) staining following etoposide treatment across glioma cell lines. We found a trend for a non-linear correlation between yH2AX staining following etoposide with susceptibility to this drug across glioma cell lines (r^2^=0.96, p=0.068 Fig **2b**). Moreover, yH2AX staining following etoposide treatment was associated with activated/cleaved caspase 3 across glioma cell lines (r^2^=0.98, p=0.0146 Fig **2c**).

**Fig 2:**
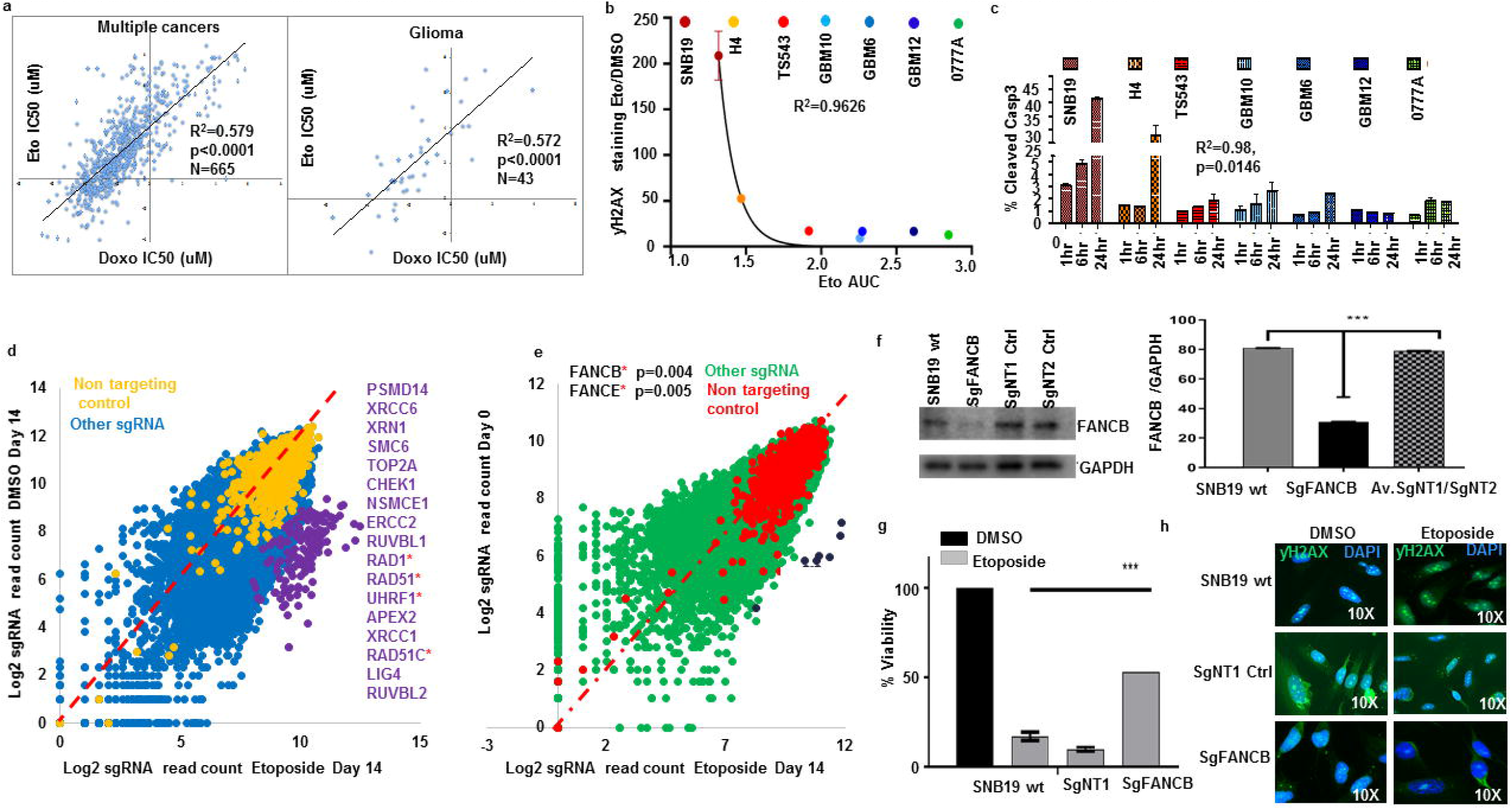
DNA damage and repair response contribute to GBM shared genetic susceptibility to TOP2 poisons. **(a)** Scatter plot for susceptibility (IC50) to etoposide and doxorubicin across human cell lines from multiple cancers (n=665, left), and the human glioma cell line subset (n=43, right) from COSMIC dataset^24^, correlation determined by Spearman’s test **(b)** Non-linear regression shows the correlation between DNA damage response (yH2AX staining) and etoposide susceptibility for different gliomas captured by the area under the curve (AUC) based on the Pearson’s correlation (Exponential growth equation). **(c)** Activated/cleaved caspase 3 was determined through flow cytometry following 1, 6 and 24 hr of etoposide 5 μM treatment. Data was normalized over DMSO treatment for each time point. Glioma cell lines ranked by etoposide susceptibility (most susceptible left, most resistant right). **(d)** Scatter plot shows the highest enriched DNA damage and repair genes enriched by etoposide compared to DMSO (p<0.001) or compared to Day 0/puromycin (p<0.01) SgRSEA enriched (Wilcoxon test) **(e)**. For **(d, e)**, the DNA damage and repair genes are ranked in order of enrichment. Genes in asterisks belong to the Fanconi anemia group of proteins. **(f)** Western blot for FANCB in the KO cells, wild type SNB19 and two clones that were edited with non-targeting control guides. Quantified against normalized GAPDH (***p<0.001). **(g)** Viability of SNB19 WT, SNB19 non-targeting control 1 (NT1), and SNB19 FANCB KO following treatment with DMSO or etoposide 5μM for 72hrs. T-test p value against FANCB vs WT SNB19 or NT1 or NT2 has (*** p<0.00001, two-tailed t-test). (h) □H2AX staining on wild type SNB19 and FANCB edited cells and the non-targeting controls treated with etoposide or DMSO, quantified in Supplementary Fig **3C**.

Next, we sought to understand which DNA damage and repair genes were enriched in either of our screens. Genes involved in DNA damage response whose KO was enriched by etoposide compared to DMSO included TOP2A (Fig **2d** and Supplementary Fig **1E**), the canonical target of TOP2 poisons, as well as SMC6 and ERCC, which are cohesin and excision repair proteins known to interact with TOP2A and TOP2B^25^. Most importantly, genes from the Fanconi anemia pathway were also present in this list, including RAD1, RAD51, RAD51C, UHRF1 (Fig **2d**). Analysis of genes involved in DNA damage and repair whose KO was selected by etoposide compared to Day 0 revealed FANCB and FANCE, which are components of core complexes of Fanconi repair machinery as the most enriched among DNA damage genes in etoposide vs Day 0 (Fig **2E**).

FANCB is a key protein within the Fanconi anemia core complex. This protein has been previously established to play a role in the DNA damage and repair pathway that is activated following treatment with several chemotherapeutics ^26^. To validate our genome-wide screen results (Fig **2e**), we performed KO of FANCB using a single guide CRISPR approach. We confirmed on-target cleavage by this sgRNA and ruled out the possibility of off-target genome editing on the locus that was predicted as the most likely off-target through a cleavage assay^27^ (Supplementary Fig **2B**). Western blot showed a decrease of FANCB protein levels in the population of FANCB KO cells edited by CRISPR (Fig **2f**), which led to acquired resistance to both etoposide (50%) (Fig **2g**) and doxorubicin (Supplementary Fig **2D**). FANCB KO also led to a decrease in yH2AX staining following etoposide treatment for 24 hrs. in contrast to an increase of this DNA damage signaling following etoposide in the control cells (Fig **2h** and Supplementary Fig **2C**). We conclude therefore that Fanconi anemia group of proteins in particular FANCB, is a major regulator of DNA damage response under TOP2 poison in gliomas.

### Expression of ribosomal proteins predict and confer glioma susceptibility to etoposide

To discover biomarkers for TOP2 poisons, we first obtained a short list of candidate genes that are implicated by being associated with and directly influencing susceptibility to this drug. To do this, we overlapped 397 genes whose loss confers etoposide resistance from our CRISPR screens (etoposide>DMSO intersection with etoposide>/Day 0, (Fig **1d**) with genes whose expression (35 glioma cell lines RNA Seq) correlates with susceptibility to etoposide (35 glioma cell lines IC50). With this, we performed differential gene expression analysis of glioma cells with etoposide susceptibility data from CCLE, using IC50<1μM as a cutoff to define susceptible lines (n=7) and IC50 >10μM (n=11) to define resistant glioma cell lines. These cutoffs were based on the rationale that systemic administration of etoposide leads to tumor concentration of 2-6 μM in human gliomas and sensitive gliomas might have a clinical response to this drug at this concentration ^16, 17, 18^. Using a p<0.01 cutoff for significance of differential gene expression (susceptible vs resistant), 9 genes (RPS18, RPS11, RPS16, RPS6, RPL35A, POLR1C, RPP25L, C10orf2 and LYRM4) out of 397 whose KO was selected by etoposide on the CRISPR screen showed higher expression on susceptible glioma cell lines compared to resistant, with 6 of these genes being ribosomal proteins. The expression of these genes on susceptible cell lines ranged from 2.9-1650 transcripts per million (TPM). Robust expression of a gene facilitates its use as a biomarker, thus we focused on RPS18, RPS11, RPS16, as these where the top 3 genes with the highest expression on susceptible cell lines among 9 selected genes (Fig **3a-b**). A gene expression analysis including all glioma cell lines (35 glioma RNA Seq) from CCLE with etoposide AUC data (35 glioma cell lines) confirmed a significant correlation between expression of these genes with etoposide susceptibility (p<0.001, Supplementary Fig **3A**). To explore whether the expression of RPS11, 16, and 18 can distinguish tumors that are susceptible to etoposide, we performed immunofluorescence staining for these markers in intracranial glioma xenografts, and found that RPS11 staining was stronger in the glioma lines MES83 and U251 (which was originated from the same human tumor as SNB19), cell lines susceptible to etoposide, intermediate expression in GBM43 which is less sensitive to this drug than the former lines, and no staining was found on GBM6 and GBM12, which exhibit resistance to this drug (Fig **3c**). To validate the implication of RPS11 in etoposide-related cell death, we edited RPS11 in SNB19 using CRISPR. RPS11 gene editing and loss was confirmed with the cleavage assay (Supplementary Fig **3B**), and a decrease of the protein by Western blot (Fig **3d**). We then investigated the contribution of RPS11 to etoposide and doxorubicin susceptibility. Viability assay following treatment with these agents showed that RPS11 KO rendered glioma cells resistant to both drugs (Fig **3e, f**).

**Fig 3:**
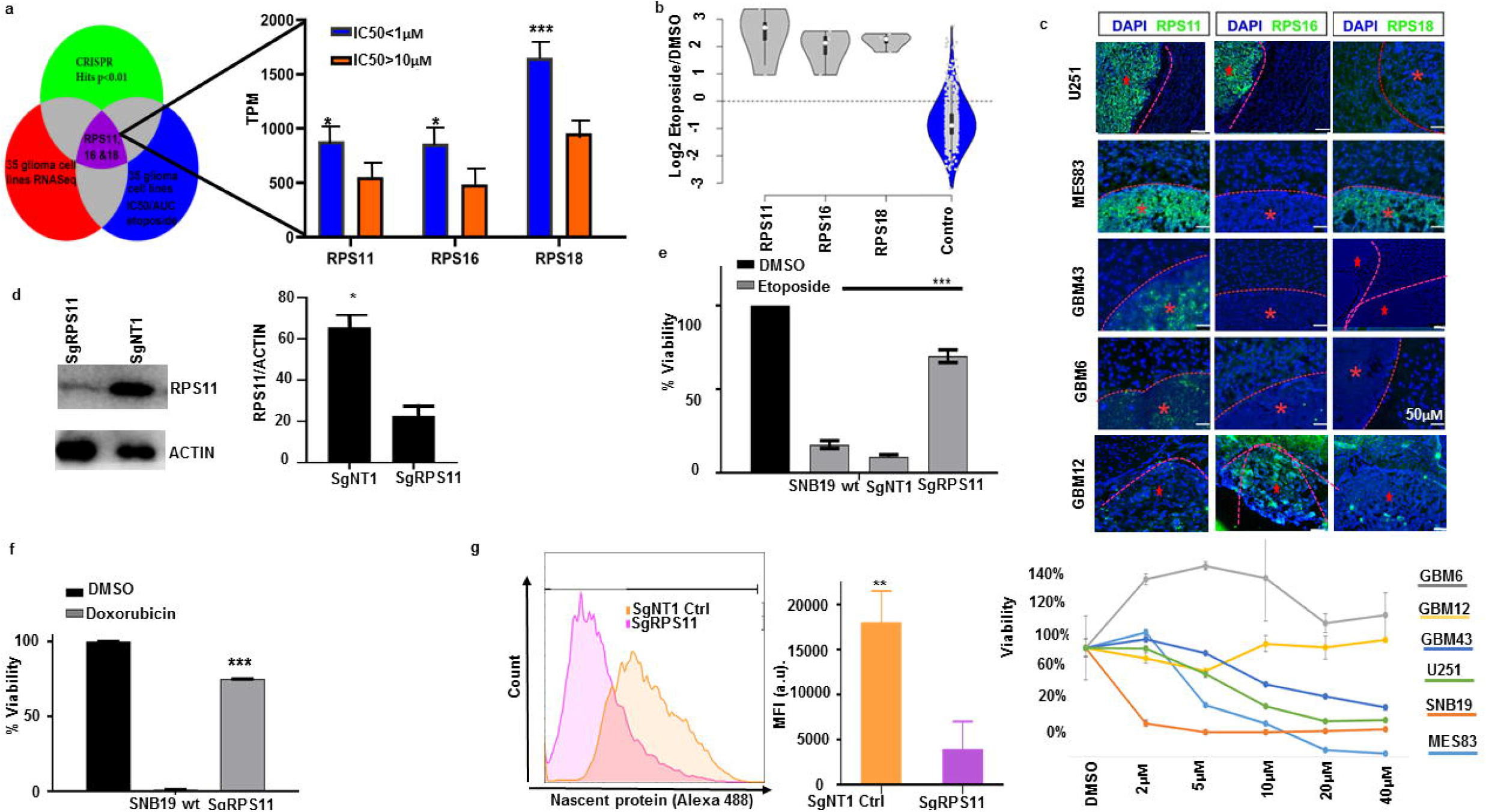
Ribosomal subunit proteins controls GBM susceptibility to TOP2 poisons and are biomarker for favorable response. **(a)** Venn diagram shows the triage of three big data sets combination (CRISPR hits p<0.01, 35 glioma cell lines RNA Seq, 35 glioma cell lines IC50/AUC) to define putative biomarker. Bar chart shows expression differences between glioma cell lines (CCLE) that are sensitive (IC50 <1μM) versus resistant (IC50 >10μM) to etoposide for RPS11 (*p=0.01), RPS16 (* p=0.0057, unpaired t-test), RPS18 (* p=0.0004, Mann Whitney test). **(b)** Violin plots show the log2 fold change enrichment of RPS11, 16, 18 in etoposide compared to DMSO and the non-targeting controls from CRISPR screen. **(c)** Immunofluorescence staining for RPS11, RPS16, RPS18 across intracranial glioma xenografts (top) with variable etoposide susceptibility determined by dose-response curves obtained *in vitro* for 72hrs (bottom). **(d)** Western blot for RPS11 on RPS11 KO SNB19 cells compared to the non-targeting controls edited SNB19 cells. Densitometry quantified (**p=0.01, unpaired t-test). **(e)** Viability assay for RPS11 KO cells (*** p=0.001), and control SNB19 cells to 5μM etoposide (left) and **(f)** 5μM doxorubicin (*** p=0.001). For **(e)**, viability data was normalized for DMSO condition for each clone. g. Histogram showing protein synthesis across RPS11 KO (purple), and non-targeting controls (orange), quantified in bar chart (**p=0.001, unpaired t-test).

We next determined the effect of RPS11 KO on translation. We labeled nascent proteins with Click-it OPP as previously described^28^, and found that RPS11 KO and the drug-resistant phenotype of these cells was associated with impaired translation (Fig **3g**). Taken together, the ribosomal protein subunits 11 controls response to TOP2 poison and is downstream of DNA damage and modulates survival from the drugs by inhibiting protein synthesis.

### Expression of ribosomal proteins is associated with cancers response to TOP2 poisons

To further explore the expression of these genes for susceptibility to TOP2 poisons, we expanded our analysis to cancers of various origins. The expression of RPS11, RPS16, RPS18 was queried in 341 cancer cell lines from multiple cancers, defining cell lines as susceptible or resistant using the same IC50 cutoff as for our analysis in gliomas. Expression of RPS11, RPS16, RPS18 remained significantly higher on susceptible cell lines from multiple cancers relative to resistant lines, and remained so for several individual cancer types (p<0.01, Fig **4a**, Supplementary List **5** and **6**). As an example, expression of RPS11, 16, 18 was significantly higher in breast cancer cell lines that were susceptible to etoposide and doxorubicin, a relevant finding considering that doxorubicin is often used to treat this disease (Fig **4b**, **4c**, Supplementary List 5).

**Fig 4:**
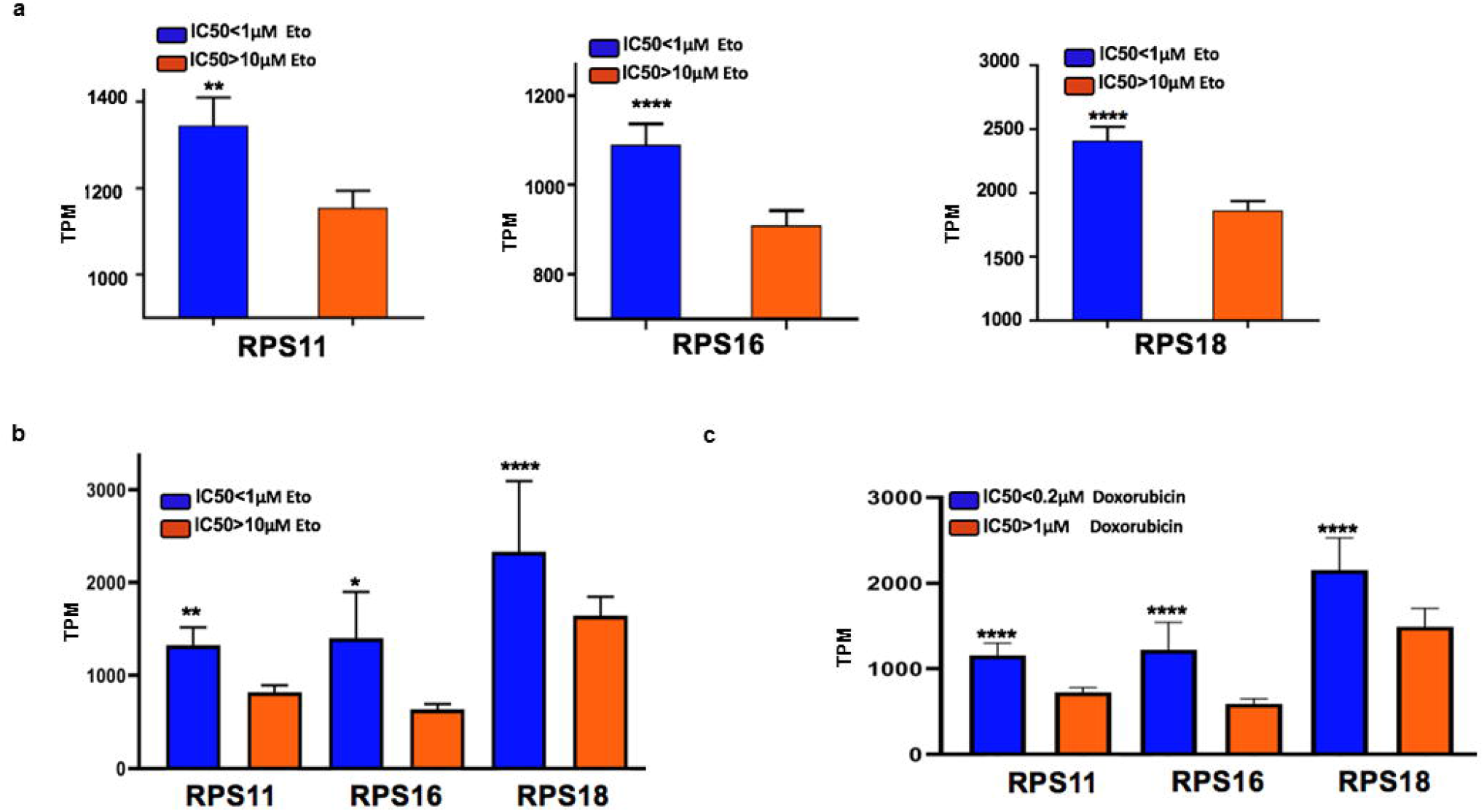
Ribosomal subunit proteins 11, 16, and 18 are influences etoposide response across cell lines for different cancers. **(a)** Bar charts shows RPS11 (** p=0.0058, Mann Whitney test), RPS16 (**** p<0.0001 Mann Whitney test), RPS18 (**** p<0.0001, Mann Whitney test) expression with IC50 (<1μM, N= 132 vs >10μM, N=209) across 341 cancer cell lines. **(b)** Bar charts shows RPS11 (**p=0.0055, two tailed t-test), RPS16 (*p=0.0161, two-tailed t-test), RPS18 (****p<0.0001, ordinary one-way Anova) expression with IC50<1μM vs >10μM etoposide for breast cancer cell lines. **(c)** Bar charts shows RPS11, RPS16 andRPS18 p<0.0001, ordinary one-way Anova expression with IC50≤0.2μM vs >1μM for breast cancer cell lines treated with doxorubicin.

### RPS11 modulates APAF1 and apoptosis during etoposide-induced translational shut-down

Previously, groups have reported chemo resistance phenotype induced by impaired ribosome biogenesis^29^. Given that translational machinery and ribosomal proteins were implicated in etoposide susceptibility, we investigated the relationship between translation, DNA damage and etoposide toxicity. First, we determined protein synthesis following treatment with this drug across multiple glioma cell lines. We found that cell lines susceptible to TOP2 poisons (SNB19, U251), showed a decrease in nascent proteins following etoposide treatment. In contrast, GBM6 and GBM12 which are resistant to TOP2, showed no decrease in nascent proteins following etoposide (Fig **5a**, Supplementary Fig **4A**, **susceptibility data on** Fig **1a** and **3c**). Given that decrease protein synthesis is associated with DNA damage across glioma cell lines, we explored the causal relationship between these two processes. RPS11 KO cells had a similar increase yH2AX foci following etoposide treatment as that seen for SNB19 wild-type or non-targeting CRISPR control cells (Fig **5b**), indicating that RPS11’s involvement in cell death is subsequent to DNA damage response activation following etoposide treatment. To investigate whether H2AX phosphorylation and DNA damage pathway activation have an effect on translation, we evaluated protein synthesis in FANCB KO cells, and observed that these suffer a decrease in nascent protein levels compared to non-targeting control cells (Fig **4b**). Based on these results, we conclude that DNA damage and H2AX phosphorylation is proximal to the effects of etoposide on translation, and therefore RPS11 might modulate viability downstream from DNA damage pathway activation.

**Fig 5:**
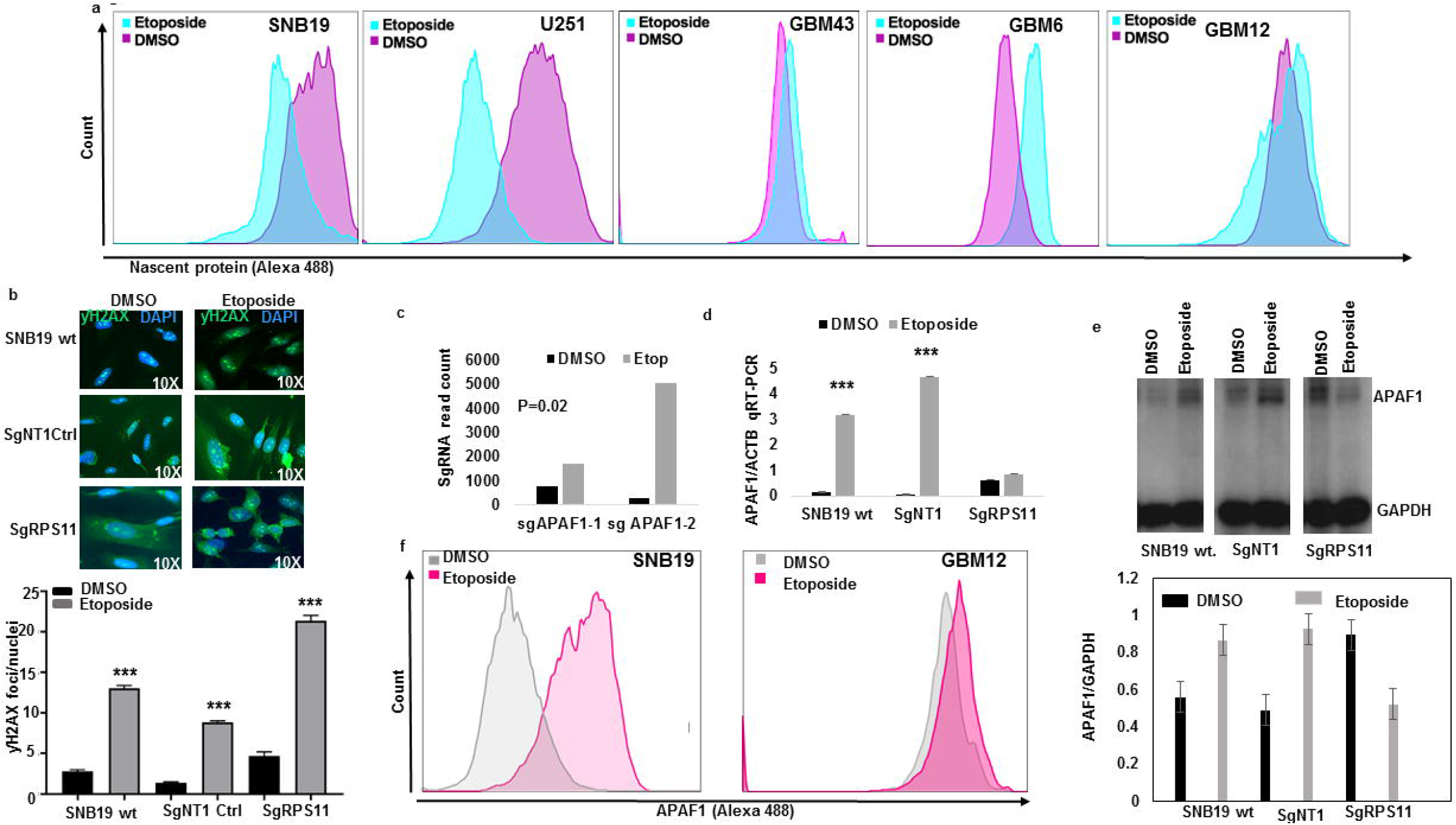
RPS11 confers susceptibility to TOP2 poisons by controlling nascent proteins and upregulating pro-apoptosome machinery APAF1. **(a)** Histogram shows the effect of 5μM etoposide treatment for 24hrs on protein synthesis across cell lines with variable degree of susceptibility (**refer to** Fig. 3c **top-bottom**). Cells are arranged in order of susceptibility (left most susceptible, intermediate susceptible, right most resistant, unpaired t-test, treated vs untreated). **(b)** □H2AX foci count on SNB19 WT cells, NT control cells or RPS11 KO with and without etoposide treatment quantified foci (below) (**** p<0.0001, zero inflated negative binomial model). **(c)** Bar plot shows the selection of two separate sgRNA for pro-apoptosome gene APAF1 in etoposide compared to DMSO from the genome-wide CRISPR screen (p=0.02). **(d)** Bar plot shows APAF1 mRNA transcript (qRT-PCR) on sgRPS11 KO, WT SnB19 and NT controls treated with and without etoposide for 24hrs (*** p=0.001, unpaired two-tailed t-test). **(e)** Western blot shows the reduction of APAF1 expression in RPS11 edited cells treated with etoposide but an increase in APAF1 in wild type and the non-targeting controls treated with etoposide (top). Bar plots show the quantification of the APAF1 expression both under etoposide and the DMSO treated cells normalized against GAPDH (bottom). **(f)** Histograms shows APAF1 expression on susceptible (SNB19) and resistant (GBM12) cell lines with and without etoposide treatment for 24hrs.

We hypothesized that RPS11 expression modulates apoptosis, as this process is triggered by DNA damage following etoposide. Whole genome CRISPR screen revealed that KO of APAF1, a key element of the apoptosis^30,31^ machinery that is activated following DNA damage was selected by etoposide compared to DMSO (Fig **5c**, Supplementary List **1B**). We then investigated the effect of RPS11 KO on APAF1 expression and how this is affected by etoposide treatment. APAF1 transcript had a significant induction following etoposide treatment in SNB19 WT and non-targeting control cells, whereas no significant expression changes were found in RPS11 KO cells (Fig **5d**). Comparison of APAF1 mRNA following etoposide between wild-type cells versus RPS11 KO suggest transcriptional modulation of this gene by RPS11. Interestingly APAF1 protein levels increased following etoposide treatment in SNB19 wild-type and non-targeting control cells, whereas etoposide treatment led to a decrease APAF1 protein in RPS11 KO cells (Fig **5e**). To investigate whether APAF1 induction by etoposide is specific to susceptible glioma lines, we compared its expression following treatment with this drug between susceptible cell line SNB19 and resistant cell line GBM12, and confirmed its induction is only seen in the former (Fig **5f**, Supplementary Fig **4C**), which we previously showed suffers H2AX phosphorylation and apoptosis following this treatment (Fig **2b-c**). These results support that RPS11 is necessary for APAF1 up-regulation following etoposide, in the context of a global shut down of protein synthesis in susceptible glioma cells at both transcript as well as protein levels.

Since APAF1 is involved at the initial phases of the apoptotic process, we explored if the pro and anti-apoptotic genes BID and BCL2^32^ downstream of APAF1 show differential expression in susceptible and resistant GBM upon TOP2 treatment. We treated a panel of GBM cell lines with and without etoposide for 24 hrs. and labelled them for BID and BCL2. These studies revealed that susceptible glioma cells show increased BID expression compared to the resistant cells (Supplementary Fig **4D-E**). Conversely, following etoposide treatment, resistant cell lines GBM6 and GBM12 showed an increase in anti-apoptotic protein BCL2 that was not observed in susceptible cell lines (Supplementary Fig **5A-G**). Taken together, our results provide mechanistic insight into why the loss of RPS11 confers resistance to TOP2 through modulation of APAF1 induction, with subsequent activation of apoptosis including BID expression and caspase 3 cleavage ^31, 32^.

## Discussion

We performed a genome scale CRISPR KO screen and combined its results with susceptibility and gene expression data from different cell lines. This approach allowed unbiased investigation of variables that account for individual tumor susceptibility to TOP2 poisons in glioma. Our CRISPR screen also revealed novel functional themes that play a role in response to this chemotherapy. In particular, we found that ribosomal subunit proteins and translation-related machinery are required to respond to these drugs.

We validated the involvement of several genes previously shown to play a role in TOP2 poison mechanism of action. These include, TOP2A, SMC6 and other genes are known to be involved in the susceptibility of cancers to TOP2 poisons^5, 6, 7, 8, 21, 22, 34^ This experiments also implicated DNA damage response and in particular, FANCB and the Fanconi anemia pathway^26^, as a key player in DNA damage response activation and susceptibility to etoposide.

Previous work implicated DNA damage response in the mechanism of action of TOP2 poisons ^5, 6^ yet to our knowledge differences in susceptibility have not been linked to this process before. Our results showed a trend for a correlation between H2AX phosphorylation following etoposide and susceptibility to this drug across glioma cell lines.

Our study provides evidence that DNA damage response activation is associated with translational modulation, and that this interaction is a major determinant of susceptibility to TOP2 poisons. These experiments indicate that DNA damage response activation (e.g. yH2AX foci) are proximal and independent of the effect of translation in susceptibility to etoposide. Indeed, KO of FANCB led to a decrease in DNA damage response activation, a decrease in protein synthesis, and acquired resistance to TOP2 poisons. On the other hand, RPS11 KO led to impairment of translation and acquired resistance to etoposide and doxorubicin, but had no significant effect on DNA damage response activation following etoposide treatment. These findings suggest that differences in DNA damage response following etoposide are independent of changes in translation in response to this drug, yet the former process influences the latter. More so, we present a novel evidence that FANCB controls nascent protein synthesis.

Our study directly implicates several ribosomal subunit proteins in response to etoposide. RPS proteins have been connected in response to chemotherapy, for example, the loss of RPS19 conferred cytoprotection to TOPI agent campotheticin ^29, 21^, previously showed RPS21, RPS27L, RPS24, RPS6KB2, RPS4Y2 and RPS4Y1 to confer susceptibility to etoposide in acute pro-myelocytic leukemia cell line (HL60).

Triggering of apoptosome machinery in the context of TOP2 poison and DNA damaging agents has also been previously reported ^30, 31^. On the other hand, the induction of apoptosis by etoposide has been linked to p53 ^29,35^. In our CRISPR screen, KO of p53 was not selected by etoposide. Yet, we show that expression of RPS11 was necessary for induction of apoptotic protein APAF1 following etoposide treatment, in the context of a robust translational shutdown that is only seen in susceptible cell lines. Given this and the fact that APAF1 KO clones were selected by our CRISPR screen, we conclude that baseline expression of RPS11 ribosomal subunit protein is necessary for induction of APAF1 and apoptosis triggering in response to etoposide, in spite of the global translational shutdown susceptible cells. Nevertheless, we acknowledge that this process is complex and might involve other mechanisms that we did not explore.

Etoposide and TOP2 poisons are highly efficacious chemotherapy agents. Yet, differences in resistance across tumors and the toxicity associated with such treatments undermines the risk/benefit ratio that this therapy can offer to individual patients ^36, 37, 38, 39, 40, 41^. To overcome this, biomarkers for etoposide and doxorubicin response are necessary, but virtually non-existent. Our work suggests that ribosomal subunits and in particular RPS11 expression might serve as predictive biomarkers for TOP2 poisons sensitivity in gliomas and across different kinds of tumors. Future prospective studies for clinical validation are necessary to establish the accuracy and clinical value of these biomarker candidates.

The use of TOP2 poisons based on individual tumor biology as opposed to histological criteria might enhance efficacy achieved by these drugs on specific patients, avoid unnecessary drug-related toxicity in patients whose tumor will not respond, and could open therapeutic options for aggressive malignancies such as gliomas. Our work sets the foundation for this precision medicine approach for the use of TOP2 poisons for gliomas, and adds to the body of evidence suggesting that the study of individual tumor biology rather than global cancer phenotype might provide more effective therapeutic interventions.

## Methods Details

### SgRNA design and lentiviral production

For loss of function screen, we used the Brunello Library that contains 70,000 sgRNA which covers the 20,000 genes in the human genome at the coverage rate of 3-4sgRNA/gene plus 10,000 sgRNA which are non-targeting controls ^12^. To prepare the library, we used the protocol as described^14^. Briefly, the HEK293T cells are grown to 70% confluence. The cells are harvested and seeded into T225 flask for 20-24hrs. The cells are mixed with Opti-MEMI reduced serum with pMD2.G −5.2μg/ml, psPAX-10.4μg/ml, Lipofectamine plus reagent and incubated with the cells for 4hrs. At the end of 4hrs the media is collected and filtered with 0.45μM filters. The virus is aliquoted and stored at −80C.

### Viral titer

To determine viral titer, 3×10^6^ of SNB19 cells are seeded into 12-well plate in 2ml. Supernatant containing virus are added at 400μl, 200μl, 100μl, 75μl,50μl, 25μl and 8μg/μl of polybrene is and spinfected at 1000g at 33°C for 2hrs. Cells then are incubated at 37°C. After 24hrs the cells are harvested and seeded at 4×10^3^ with puromycin for 96hrs with a well containing cells that were not transduced with any virus. After 96hrs the titre glo is used to determine cell viability at MOI 21%. At the multiplicity of infection (MOI) of 21% we are able to infect 1 sgRNA/cell.

### Large scale cell culture and expansion

To perform the CRISPR screening, SNB19 cells were expanded to 500million and then spinfected with 70,000sgRNA. After spinfection, the cells are selected with 0.6μg/ml of puromycin for 4days. This selection is aimed at the cells that have been rightly integrated with the sgRNA that incorporates the puromycin cassette into their genome. We achieved an MOI of 21% in two independent screens. At the end of day 4, about 150million cells survived the selection. We used 50 million of selected cells for the extraction of genomic DNA. The base sgRNA representation is obtained by amplification of the sgRNA with unique barcoded primers. The remaining 100million cells were expanded for 2days, once cells grew to 200 million. 100 million of cells were treated with etoposide at concentration of 5 μM for 14 days, and the remaining 100 million were treated with DMSO for 14 days and served as control. After 14days, the cells were harvested, the gDNA extracted, and the sgRNA amplified with another unique barcoded primer.

### DNA extraction and PCR amplification of pooled sgRNA

Briefly, the genomic DNA (gDNA) were extracted with the Zymo Research Quick-DNA midiprep plus kit (Cat No: D4075). gDNA was further cleaned by precipitation with 100% ethanol with 1/10 volume 3M sodium acetate, PH 5.2 and 1:40 glycogen co-precipitant (Invitrogen Cat No: AM9515). The gDNA concentration were measured by Nano drop 2000 (Thermo Scientific). The PCR were set up as described^14^. The sgRNA library, puromycin, DMSO and etoposide selected guide RNA were all barcoded with unique primers as previously described^14^.

### Next Generation Sequencing

The sgRNAs were pooled together and sequenced in a Next generation sequencer (Next Seq) at 300 million reads for the four sgRNA pool aiming at 1,000reads/sgRNA. The samples were sequenced according to the Illumina user manual with 80 cycles of read 1 (forward) and 8 cycles of index 1^14^. 20% PhiX were added on the Next Seq to improve library diversity and aiming for a coverage of >1000reads per SgRNA in the library.

### CRISPR screen data analysis

All data analysis was performed with the bioinformatics tool CRISPR Analyzer^45^. Briefly, the sequence reads obtained from Next Seq were aligned with human genome in quality assessment to determine the percentage that maps to the total human genome. To set up the analysis, the sgRNA reads (library, puromycin, DMSO and etoposide) replicates were loaded unto the software. The sgRNA that does not meet a read count of 20 is removed. Hit calling from the CRISPR screen was done based on sgRSEA enriched, p<0.01 was used for significance based on Wilcoxon test.

### Gene Ontology

We used DAVID^42, 43^ and analyzed for the biological pathways that were enriched for etoposide and genes that controls glioma susceptibility to TOP2.

### Immunofluorescence

Northwestern University institutional animal care facility (IACUC) approved the animal experiments. GBM patient-derived xenograft (PDX) lines, MES83, U251, GBM6, GBM12 and GBM43, were all implanted into brain of nude mice using stereotactic device and following institutional animal care facility protocols. Once tumor implanted (4-6weeks), we sacrificed the animal and fixed the brain with 4% PFA. Using sucrose gradient, we dehydrated the tissue and mounted with OCT. Tissue section were cut at 5μM. We washed the tissue in PBS-tween20, and incubated with anti-RPS11, 16 and18 (1:100) (Supplementary List **4**) overnight at room temperature. Tissue were blocked in 3% BSA in PBS and incubated for 2hrs. Using anti-rabbit Alexa 488 and DAPI mounting media, we stained the proteins and obtained images on confocal microscope.

### Single gene editing

To edit FANCB and RPS11, we used single guide RNAs that were enriched for both genes as well as the non-targeting controls (Supplementary List 3). Briefly, these guides were synthesized by Synthego and following the protocol, we prepared the ribonucleoprotein complexes by mixing the guides (180pmol) with recombinant Cas9 protein (Synthego) 20pmol in 1:2 ratio. The complexes were allowed to form at room temp for 15mins, and then 125μL of Opti-MEM I reduced serum medium and 5μL of lipofectamine Cas9 plus reagent were then added. Both, the cells and the formed ribonucleoprotein complexes, were seeded at the same time with 150,000 SNB19 cells in a T25 flask. The cells were incubated for 4days. After 4 days, the cells were harvested, and downstream analysis were performed to prove the editing of the genes.

### T7E1 cleavage assay

To confirm the efficiency of the edit, we extracted the gDNA from the edited cells as described from Gene Art (Cat No: A24372) and then using primers listed in (Supplementary List **3**) for the on-target and the off targets of FANCB and RPS11 respectively and amplified them by PCR. The PCR cycle used has been described in Gene Art (Cat no: A24372). The amplified bands were gel extracted and hybridized as described in Gene Art (Cat no: A24372). Subsequently, we incubated the hybridized amplicon with T7E1 (NEB: M0302). The cleaved bands were resolved on 2% agarose gel.

### Western blot

To confirm the loss of protein expression of the gene of interest following editing, we extracted the proteins using M-PER (Thermoscientific: 78501) and cocktail of phosphatase and protease inhibitors. The cells were lysed using water bath ultrasonicator for 4mins. Cell lysate were cleared by centrifugation. We measured the concentration of protein in lysates. Denatured lysates were loaded into 4-20% Trisglycine gels (Novex) and separated at 180V for 2hrs. The gels were transferred unto a PVDF membrane by semi-dry blotting for 1hr. We blocked the membrane in 5% non-fat milk TBST buffer for 30mins and incubated with primary antibodies RPS11 (1:500),FANCB (1:500),GAPDH or ACTB (1:1500) in 5% BSA respectively over night shaking at 4°C. Primary antibodies were removed and we added the secondary polyclonal HRP (1:20,000) in TBST and incubated shaking for 2hrs at room temp. The membrane were washed 6X in TBST and then developed with ECL (Cat No: 1705061) and band imaged on a Bio-Rad Chemi-doc imaging system.

### Viability assay

The edited cells (FANCB, RPS11, non-targeting control and wild type unedited SNB19) were seeded at 4,000 cells/well in a 96 well plate and treated them with 5μM etoposide and doxorubicin or DMSO for 72hrs. For GBM PDX lines, we seeded them as well at 4,000cells/well in a 96 well and then added etoposide at a range of 2-40μMfor 72hrs. Titre glo was added following incubation with drugs. (Cat No: G7572) and the viability of cells was analyzed 5 min later by measuring the luminescence. We normalized the intensity against DMSO treated cells of each cell line or PDX or the edited cell and then determined the survival. Pictures of these cells were also taken as shown in the source data figures.

### Click–it Plus OPP Assay, Apoptosis Assay and Flow Cytometry

To determine if nascent protein synthesis is impaired upon editing of RPS11 and FANCB. We seeded edited cells SgRPS11, SgFANCB, wild type SNB19 and the non-targeting controls in a 96 well plate with black covers overnight at 4,000cells/well. To determine if etoposide impacted nascent protein synthesis and apoptosis on the GBM PDX lines, we treated them with DMSO or etoposide 5μM for 24hrs. Following the protocol from Life technologies (Cat No: C10456), we added the Click-it OPP (1:1000), or Caspase 3/7 (Cat no: C10427) or with antibodies against BID,BCL2 or APAF1 for 30mins.After washing cells were fixed and primary antibodies were detected secondary antibodies conjugated to Alexa Flour 488. The fluorophore intensity was measured by flow cytometry (LSR Fortessa 1 analyzer). As a complementary approach, the labelled cells were also imaged in a fluorescent microscope (Nikon Ti2 Widefield).

### qRT-PCR

Wild type and the sgRPS11 edited SNB19 cells as well as the non-targeting controls cells were treated with and without etoposide (5μM) for 24hrs. The total RNA was extracted using Zymo Research kit (Direct–zol RNA Miniprep Plus, Cat no: R2070). The quality of the RNA was determined by RNA pico bioanalyzer measuring the 18S and 28S ribosome. Using the superscript III first strand synthesis system (Cat no: 18080-051), we generated cDNA and performed qPCR with APAF1 and ACTB primers in triplicates and fold change of expression of APAF1 were normalized against actin B (ACTB).

### DNA damage assay

For the analysis of DNA damage cells were seeded at 4,000/well together with wild type cells and the non-targeting control edited cells. Cells were treated with 5μM etoposide for 24 hrs. The cells were harvested and then using the protocol from BD Science (material no: 560477), the cells were fixed with 200μL of 4% PFA for 10mins, blocked in 10% BSA for 2hrs at room temperature, washed with PBS, and then 1:10 H2A.X antibody phospho S139 (ab11174) were added and incubated for overnight. After washing, the primary antibody was detected by goat anti-mouse antibody conjugated to Alexa Flour 488(Thermofisher: #A-11001). DAPI nuclear stain in mounting media was used to counterstain nucleus. We obtained images of foci of gamma H2AX and using Nikon element imaging software (NIS-element), we counted the foci inside the nucleus.

### GBM patient derived xenograft culture

The patients derived xenografts GBM 12, GBM6, GBM83, and GBM43 were used in this study. Briefly, all the GBM PDX cells were all authenticated, they were cultured in 1% FBS in DMEM media. SNB19 were grown in 10% FBS in MEM media containing, essential amino acids, sodium pyruvate and 1% glutamine. U251 were grown in 10% FBS DMEM media. The cells were all grown to 80% confluency and then used for downstream analysis.

### Statistical analysis

Briefly, the CRISPR analysis were all performed with CRISR Analyzer^45^ which contains 8 statistical analysis for hit calling. All our experiments were performed in at least two independent experiments with multiple replicates. All bar charts in the manuscript were built with Graph Pad prism software 8 (San Diego, CA, USA). The statistical analysis performed for each figure are listed in the figure of the accompanying figures.

### Reagents used

All reagents used in this work are listed in Supplementary List **4**

## Supporting information

Supplemental List 1A

Supplemental List 1B

Supplemental List 2A

Supplemental List 2B

Supplemental List 3

Supplemental List 4

Supplemental List 5

Supplemental List 6

Supplemental List 7

## Author’s contribution

AS and CUA conceived the idea of this project. CUA, AM, LC performed the genome scale CRISPR screen. JW and CUA performed the screen analysis. MB and CUA performed the combinatorial analysis to predict biomarkers. CUA performed the single gene editing and all validations. ML contributed to organizing and curating the largescale data used in this work. CUA, LW and DZ performed western blots. EF contributed in cell culture. DY and CUA performed the immunohistochemistry, EBG and CP transduced lentivirus with RPS11 overexpression vector. MZ and CUA performed the flow cytometry analysis. SK contributed to the CRISPR screen. IB contributed reagents and corrected the manuscript. CUA and RM performed the DNA damage analysis on edited cells and GBM cell lines respectively. BG, KMB, DMS performed oversaw statistical analysis. AS, PH, AH supervised the project. CUA, MZ and AS prepared figures. CUA and AS wrote the manuscript.

## Acknowledgement

This work was funded by 5DP5OD021356-04 (AS), P50CA221747 SPORE for Translational Approaches to Brain Cancer (AS), and Developmental funds from The Robert H Lurie NCI Cancer Center Support Grant #P30CA060553 (AS). We thank Dr Ichiro Nakano (University of Alabama), Dr. Charles David James (Northwestern University) and Dr. Shi-Yuan Cheng (Northwestern University), for the kind gift of the GBM xenografts. We thank Dr. Peng Zhang (Northwestern University) for technical support. We thank Synthego CA, USA for the gifts of RPS11 sgRNA guides. □H2AX imaging work was performed at the Northwestern University Center for Advanced Microscopy generously supported by NCI CCSG P30 CA060553 awarded to the Robert H Lurie Comprehensive Cancer Center. BG was supported by the NVKP 16-1-2016-0037 grant.

## Declaration of Interest

The authors declare conflict of interest in that a patent application has been submitted in regard to the findings here in this article.

## Corresponding author

Adam M Sonabend MD

