## Supplementary Figure & text for "Interaction between DNA damage response, translation and apoptosome determines cancer susceptibility to TOP2 poisons"

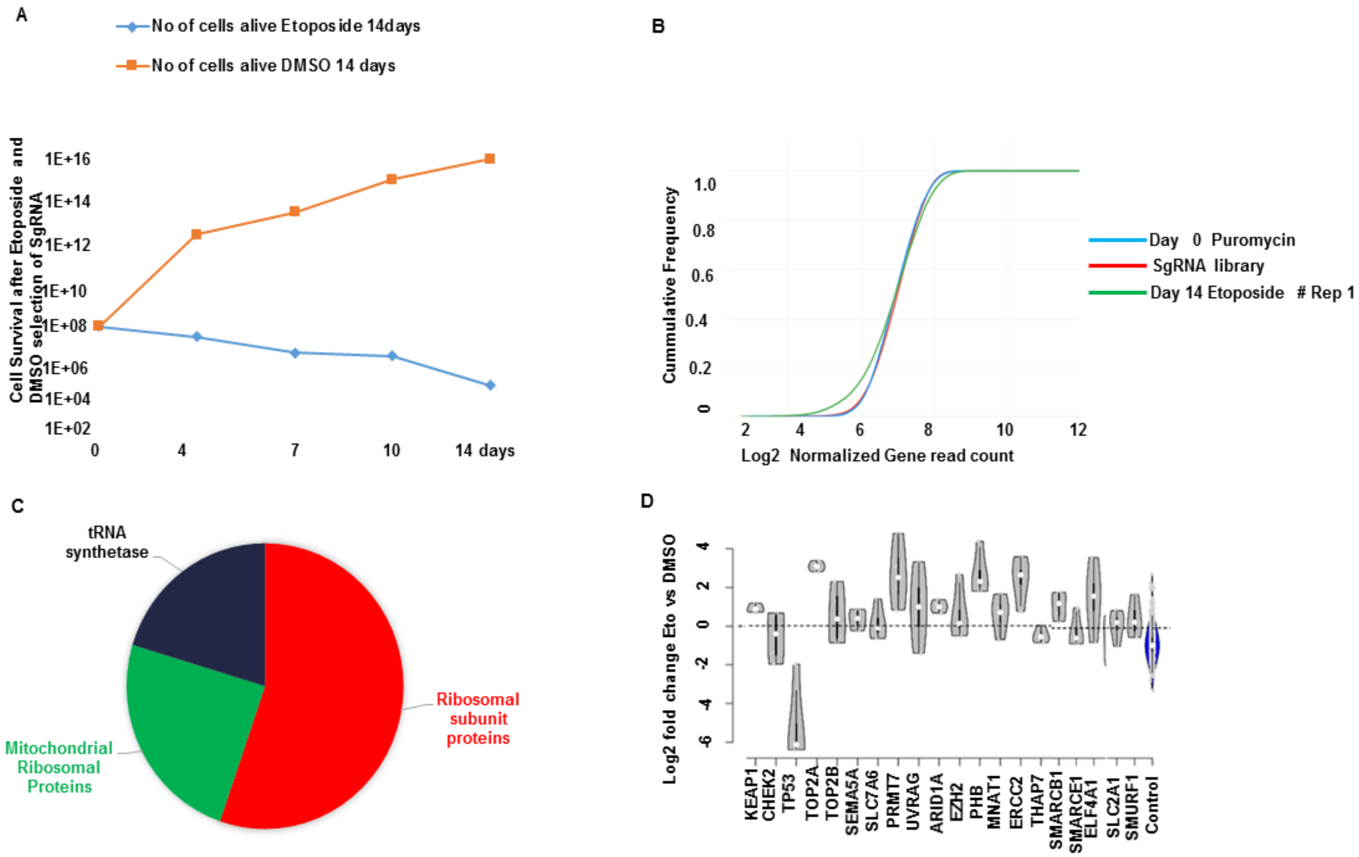


**Supplementary Fig 1: CRISPR screen cells survival, sgRNA reads and gene ontology of enriched themes (Related to Fig 1).**

**(A)** The chart shows cell counts while undergoing treatment with etoposide or DMSO for 14 days from the CRISPR KO screen experiment. **(B)** Comparison of cumulative frequency of sgRNA for the Brunello library plasmid used for generating the lentiviral particles (sgRNA library red), puromycin selection (Day 0 blue), and etoposide treated cells replicates 1 (Day 14 Etoposide # Rep 1). **(C)** Pie chart shows that within the gene ontology themes related to the translation machinery, the most genes were related to ribosomal subunit proteins (n=70), followed by mitochondrial ribosomal proteins (n=30) and tRNA synthetase (n=25). **(D)** Violin plot shows genes (KEAP1 p=0.013, TOP2A p=0.00039, PRMT7 p=0.00093, UVRAG p=0.002971, ARID1A p=0.05, PHB p=9.84E-05, MNAT1 p=0.02, ERCC2 p=0.000683, SMARCB1 p=0.0226, ELF4A1 p=0.02, SLC2A1 p<0.005, SMURF1 p=0.01) known to contribute to etoposide susceptibility were enriched in etoposide (day 14) compared to DMSO (day 14). However, TP53, CHEK2, SLC7A6, THAP7 and SMARCE1 were not enriched.

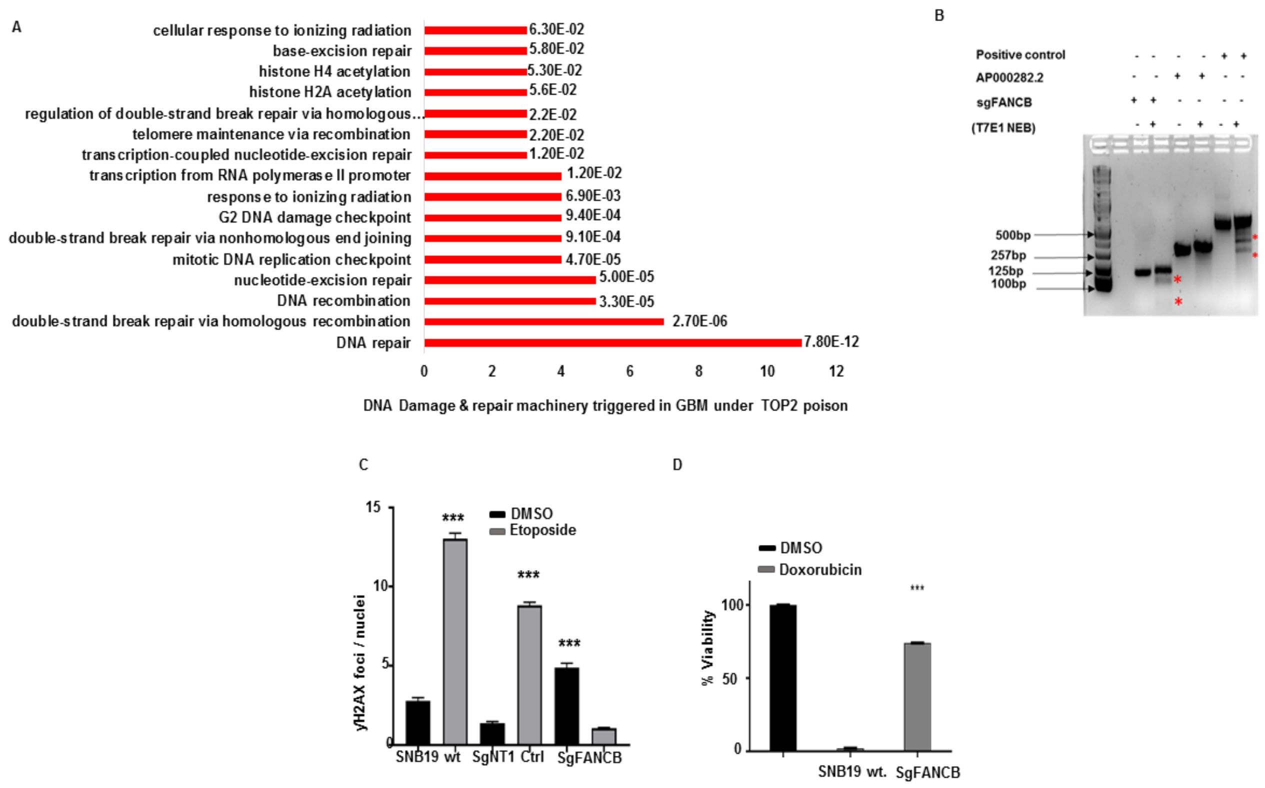


**Supplementary Fig 2: Loss of FANCB confers resistance to etoposide, doxorubicin and DNA damage lesion and repair processes at play under etoposide in GBM (Related to Fig 2).**

**(A)** List of distinct DNA damage and repair processes implicated by genes whose KO was selected by etoposide. **(B)** Cleavage assay editing in the on-target FANCB but no off-target cleavage on the AP000282.2, including positive control cells edited at other sites with TALEN. **(C)** Graph shows quantified count of γH2AX foci on SNB19 wild type, SgNT1 control and SgFANCB treated with and without etoposide (****$p<0.0001$, zero-inflated negative binomial model). **(D)** The viability assay quantified shows significant survival of FANCB edited cells under 5µM doxorubicin treatment for 72hrs compared to wild type cells (***$p=0.001$).

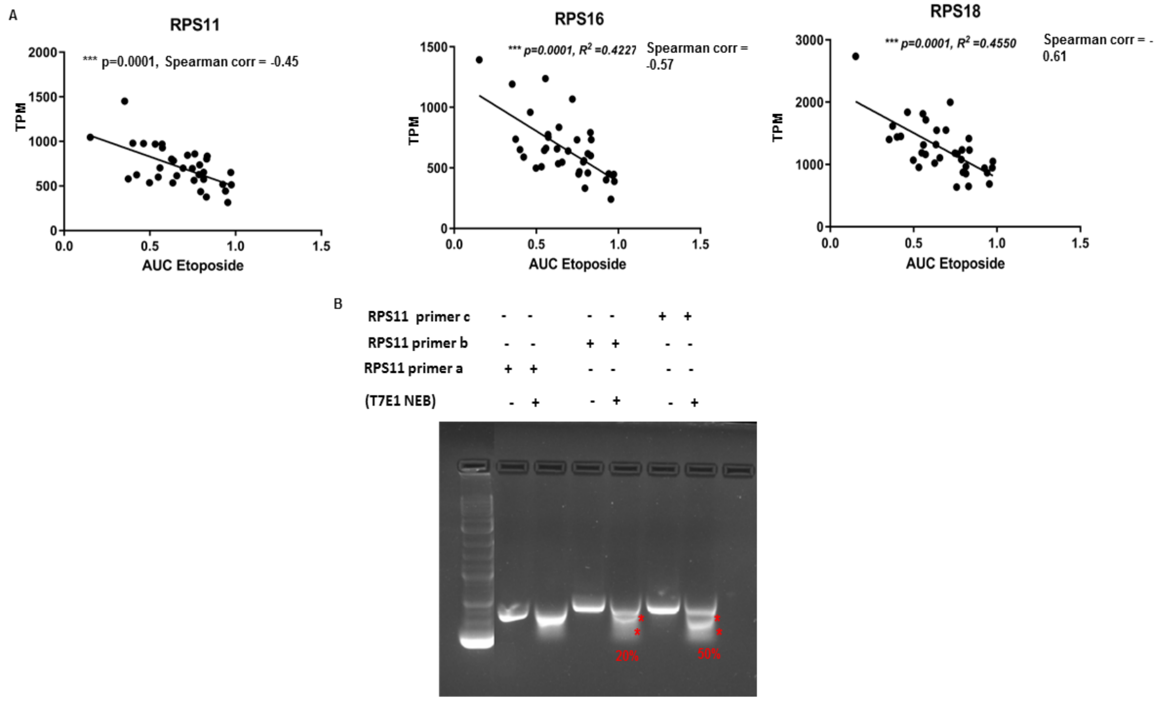


**Supplementary Fig 3: Expression of Ribosomal subunit proteins correlates with IC50 and AUC etoposide for glioma and many cancers, sgRPS11 edits efficiently (Related to Fig 3).**

**(A)** The figures show the correlation of the predicted biomarkers RPS11 (***p=0.0001), RPS16 (*** p=0.0001), RPS18 (***p=0.0001), TPM vs their AUC etoposide (inverse spearman's correlation) across 36 glioma CCLE cell lines. **(B)** The gel shows cleavage assay of sgRPS11 edited cells with 50% cleavage at target sites. The cleavage was confirmed with three independent primer sets.

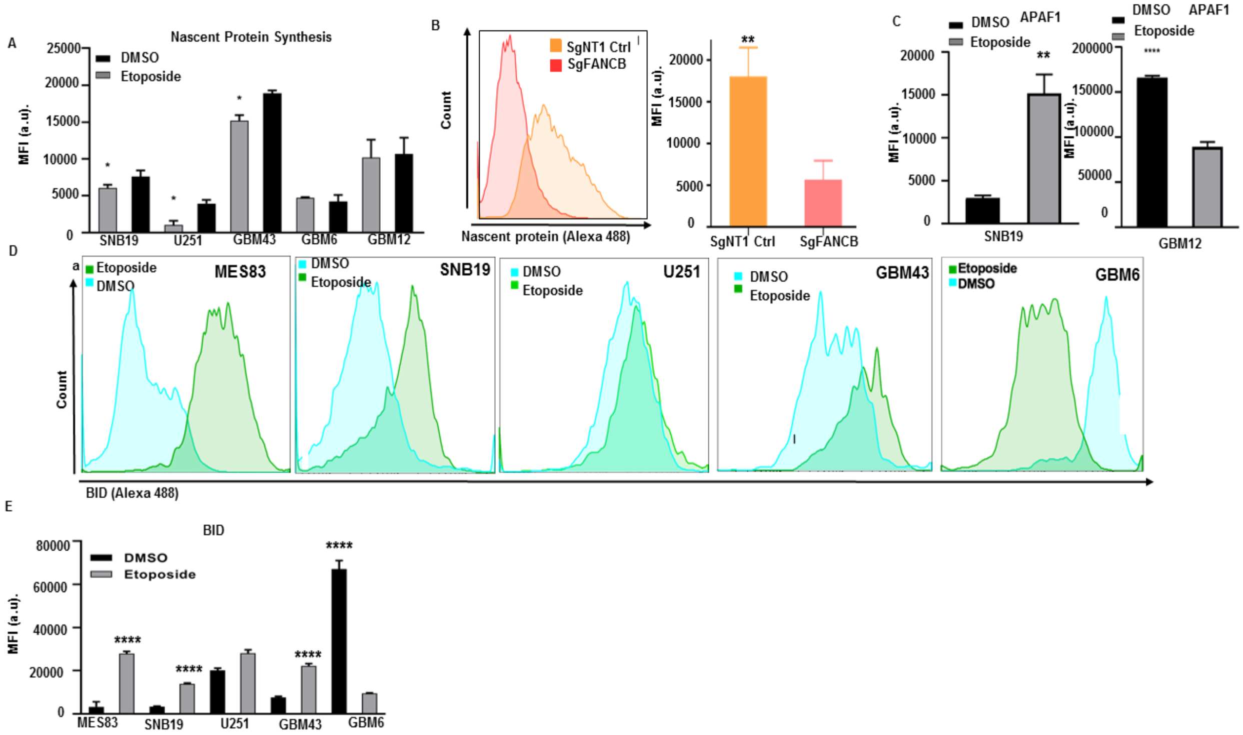


**Supplementary Fig 4: BID shows differential expression in GBM susceptibility to TOP2 poison committed to apoptosis (Related to Fig 5).**

**(A)** Bar plots shows median fluorescence intensity of nascent protein synthesis across GBM with and without etoposide for 24hrs. **(B)** Histograms shows FANCB KO (red), SgNT1 (orange) labelled with Click-it OPP to determine protein synthesis (left), which was quantified in bar plots (right) (***p=0.0001, unpaired t-test). **(C)** Bar plot shows median fluorescence intensity of APAF1 between SNB19 (right **p<0.0098, unpaired t-test with Welch's correction) and GBM12 (**** p<0.0001, unpaired t-test) treated with and without etoposide for 24hrs. **(D)** Histogram shows BID expression in glioma cell lines following etoposide, in which cell lines were ranked based on susceptibility to this drug (left most susceptible and right most resistant). The median fluorescence intensity for BID expression quantified **(E)** (**** p<0.0001, unpaired t-test).

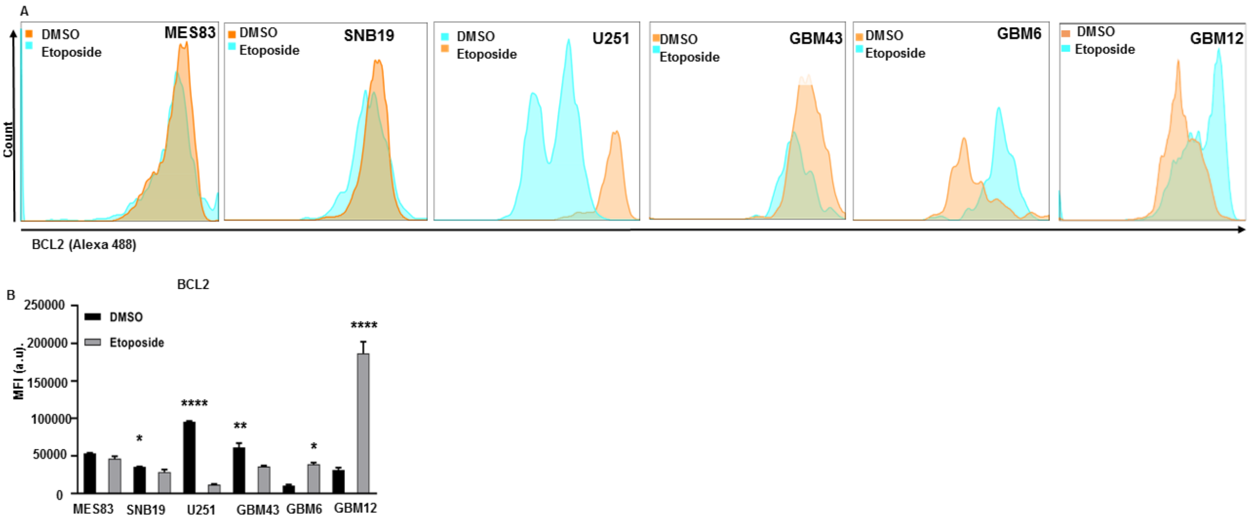


**Supplementary Fig 5: BCL2 shows differential expression in GBM susceptibility to TOP2 poison committed to apoptosis.**

**(A)** Histogram shows BCL2 expression in GBM cell lines treated by etoposide for 24hrs arranged from susceptible (left) to resistant (right) **(B)** The median fluorescence intensity quantified (* p=0.02, **p=0.0014, ***p=0.0001, **** p=0.0001, unpaired t-test).

**Supplementary List 1A: Statistics of the CRISPR screen replication correlations.**

**Supplementary List 1B: Genes that are enriched p<0.01 in (etoposide vs DMSO) and (etoposide vs Puromycin (Day 0)) and the genes that overlapped between them.**

**Supplementary List 2A: Gene ontology of the enriched genes from the CRISPR screen (Etoposide vs DMSO) p<0.001.**

**Supplementary List 2B: DNA damage repair genes enriched in the screen (Etoposide vs DMSO) p<0.001 [20].**

**Supplementary List 3: Guide RNAs used in the single gene editing, on target and off target positions and the PCR primers used in this study.**

**Supplementary List 4: Reagents used in this study.**

**Supplementary List 5: Analysis RPS11, 16, and 18 expression in 341 cancers that are treated with TOP2 poison.**

**Supplementary List 6: Analysis of cancers (B-cell lymphoma, Burkitt's lymphoma, Hodgkin's lymphoma, Lung non-small cell carcinoma, Lung small cell carcinoma, Neuroblastoma and Ovarian cancers) treated with TOP2 that RPS11, 16 and 18 influences response to it (marked in red).**

**Supplementary List 7: CRISPR Next Seq Read count**
